# Structure of the Quenched Cyanobacterial OCP-Phycobilisome Complex

**DOI:** 10.1101/2021.11.15.468719

**Authors:** Maria Agustina Dominguez-Martin, Paul V. Sauer, Markus Sutter, Henning Kirst, David Bina, Basil J. Greber, Eva Nogales, Tomáš Polívka, Cheryl A. Kerfeld

## Abstract

Photoprotection is an essential mechanism in photosynthetic organisms to balance the harvesting of light energy against the risks of photodamage. In cyanobacteria, photoprotective non-photochemical quenching relies on the interaction between a photoreceptor, the Orange Carotenoid Protein (OCP), and the antenna, the phycobilisome (PBS). Here we report the first structure of the OCP-PBS complex at 2.7 Å overall resolution obtained by cryo-electron microscopy. The structure shows that the 6.2 MDa PBS is quenched by four 34 kDa OCP organized as two dimers. The complex also reveals that the structure of the active form of the OCP is drastically different than its resting, non-quenching form, with an ∼60 Å displacement of its regulatory domain. These results provide a high-resolution blueprint of the structural basis of the protective quenching of excess excitation energy that enables cyanobacteria to harvest light energy and fix CO_2_ across environmentally diverse and dynamic surface of our planet.

---

All photosynthetic organisms require photoprotective mechanisms that are activated when their light harvesting antennas absorb excess light-energy. The photoprotective state is elicited not only under high light, but also in the context of drought or temperature stress. In light harvesting complexes the rapid conversion of excess excitation energy into heat before it causes harm is known as non-photochemical quenching (NPQ)^1^. NPQ is also wasteful. In cyanobacteria, organisms ancestral to the chloroplast, 60-80% of captured light energy is lost to NPQ^2^. In most cyanobacteria, NPQ is triggered by a water-soluble photoreceptor that binds a single carotenoid molecule, the Orange Carotenoid Protein (OCP)^3-6^. Absorption of blue-green light by the resting, orange form of the protein, OCP°, converts it into the quenching red form, OCP^R^ that binds to the phycobilisome (PBS) antenna^7-10^. The structure of the OCP^R^ and how it binds to the PBS have evaded elucidation resulting in ongoing debates concerning the mechanism of OCP-induced energy dissipation. Here we report the structure of the OCP-PBS complex from the model organism *Synechocystis* PCC 6803 at an overall 2.7 Å resolution. We show that the PBS is, in contrast to all previous predictions, quenched by two OCP dimers at two distinct sites of the PBS core. Moreover, our structure reveals an unanticipated drastic domain displacement that takes place when OCP° is converted to OCP^R^. We also show that quenching occurs via energy transfer and that the 6.2 MDa PBS with its 396 bilin molecules is efficiently quenched by four 34 kDa OCPs. Collectively these data constitute the first structure of a quenched light harvesting antenna and provide a blueprint for the understanding and engineering the photoprotective state of the cyanobacterial PBS.

## Overview of the structure of the OCP-PBS complex and comparison to the PBS in the light harvesting state

In order to understand the structural basis of OCP-induced NPQ of the cyanobacterial PBS, we have determined the structure of the intact quenched OCP-PBS complex from *Synechocystis* PCC 6803 at an overall resolution of 2.7 Å by cryo-electron microscopy (cryo-EM), with individual domains resolved at 2.1 Å (rods) to 2.5 - 4 Å (core regions and OCP) (Fig. 1 a, Extended Data Fig. 1 and 2, Extended Data Table 1 and Methods). In addition to the PBS (consisting of 320 protein chains and 396 bilins) we resolved four OCP chains, each binding a canthaxanthin (CAN) carotenoid (Fig. 1 d) summing to a total molecular weight of 6.3 MDa. For analysis of the *Synechocystis* PCC 6803 PBS architecture in the light harvesting state we refer to our accompanying study^11^. The quenched state requires a specific PBS configuration, referred to as the up-up conformation in^11^ (Fig. 1 a). In contrast to previous biochemical and spectroscopic studies that suggested that the PBS has one or two OCP binding sites^8,12^, our cryo-EM structure of the OCP-PBS complex reveals that four OCP^R^, arranged as two dimers, are bound to the core of the light-harvesting antenna (Top, hereafter T, and Bottom, hereafter B, cylinders) (Fig. 1 a-c) in the fully quenched state.

**Figure 1:**
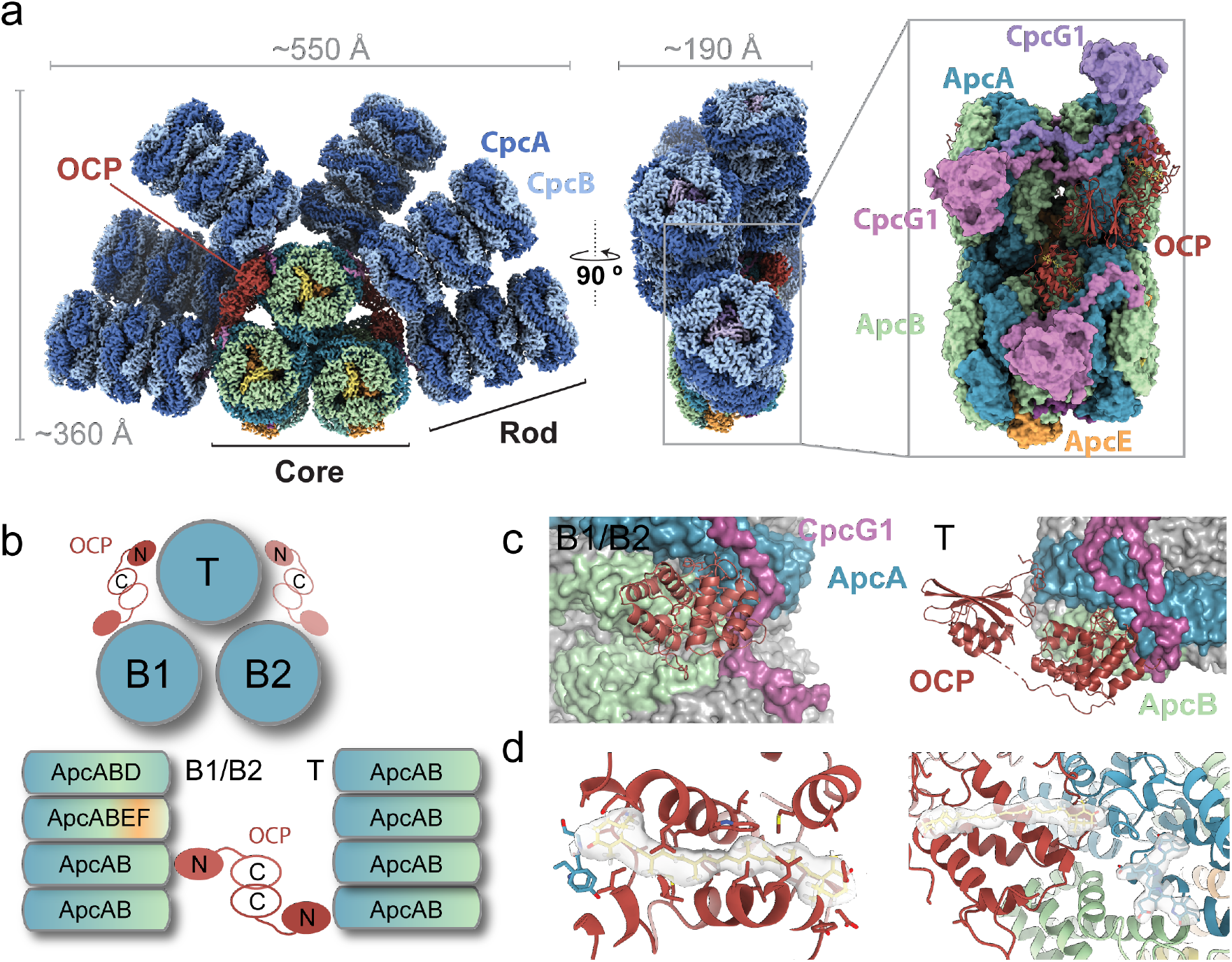
Overview of the OCP^R^-PBS structure. **a**, Composite cryo-EM map of the complete OCP^R^-PBS complex, front and side view. Individual maps and the consensus map can be found in the Extended Data Fig. 1. Inset showing a side-view in surface representation, omitting the rods. **b**, Schematic of the OCP^R^-PBS core complex. **c**, Overview of the OCP bound to the PBS in B1/B2 cylinder (Bottom cylinders) and T cylinder (Top cylinder). The PBS proteins that interact with OCP colored. **d**, Close-up views showing the density for the carotenoid and the atomic models for the carotenoid and its closest bilin.

The up-up conformation of PBS in the light harvesting state (no bound OCP^R^)^11^ and in the quenched (four bound OCP^R^) state closely superimpose (Extended Data Fig. 3). The binding of the four OCP^R^ to the core results in a slight structural perturbation (overall rmsd of 0.44 Å over 12,716 aligned core Cα atoms). Given the multiple conformations of the PBS in its light harvesting state^11^ each distinguished by different position of the top rod (denoted therefore as the mobile rod, see Extended Data Fig. 3), it is notable that the binding site of OCP is situated right below the attachment of the mobile rod to the core. Therefore, when the mobile rod is in the down position^11^, it completely occludes the OCP binding site. We do not observe any partially quenched states (e.g. consisting of only one OCP-dimer bound to the PBS), possibly because our sample contained a large excess of OCP to PBS (20:1) to favor the formation of the fully quenched complex (Extended Data Fig. 4 c, d). Our model suggests that while the mobile rod of the unoccupied side of a partially quenched PBS complexes could move freely between its up and down states the fully quenched state requires the up-up PBS conformation in order to have all four OCP^R^ binding sites accessible. Recently, the in situ higher order organization of the PBS was described^13^; it is made up of closely appressed PBS in the up-up state.

Even in this compact arrangement, the OCP binding dimer sites are accessible, suggestive of the potential for these light harvesting arrays to be rapidly quenched by the OCP (Extended Data Fig. 5).

## The Structure of the OCP^R^

Our OCP-PBS structure also reveals the structure of OCP^R^, the red active form of OCP (Fig. 2). Structurally, the inactive OCP° is composed of two domains, an N-terminal domain (NTD) with an α-helical fold unique to cyanobacteria and a mixed α/β C-terminal domain (CTD) joined by a flexible linker, a single carotenoid molecule that spans both domains^6^ (Fig. 2 a). However, the OCP^R^ bound to the PBS shows that the NTD and CTD are completely separated and only connected by an extension of the interdomain linker (Fig. 2 a, b). In addition to the elongated structure and the complete translocation of the carotenoid into the NTD consistent with previous findings^9,14^, our structure shows that in OCP^R^ the CTD has rotated ∼220° around the NTD, resulting in a net translation of about 60 Å for the center of mass of the CTD (Fig. 2 b, Supplementary video 1). This translation is slightly different for the T vs B cylinder bound state (61 Å for T, 63 Å for B). The translocation of the carotenoid from a position shared between the N- and C-terminal domains completely into the NTD along with the detachment of the CTD exposes the β1 surface, which is buried between both domains in OCP°, and thus allows it to interact with the binding site on the PBS (Fig. 2 a).

**Figure 2:**
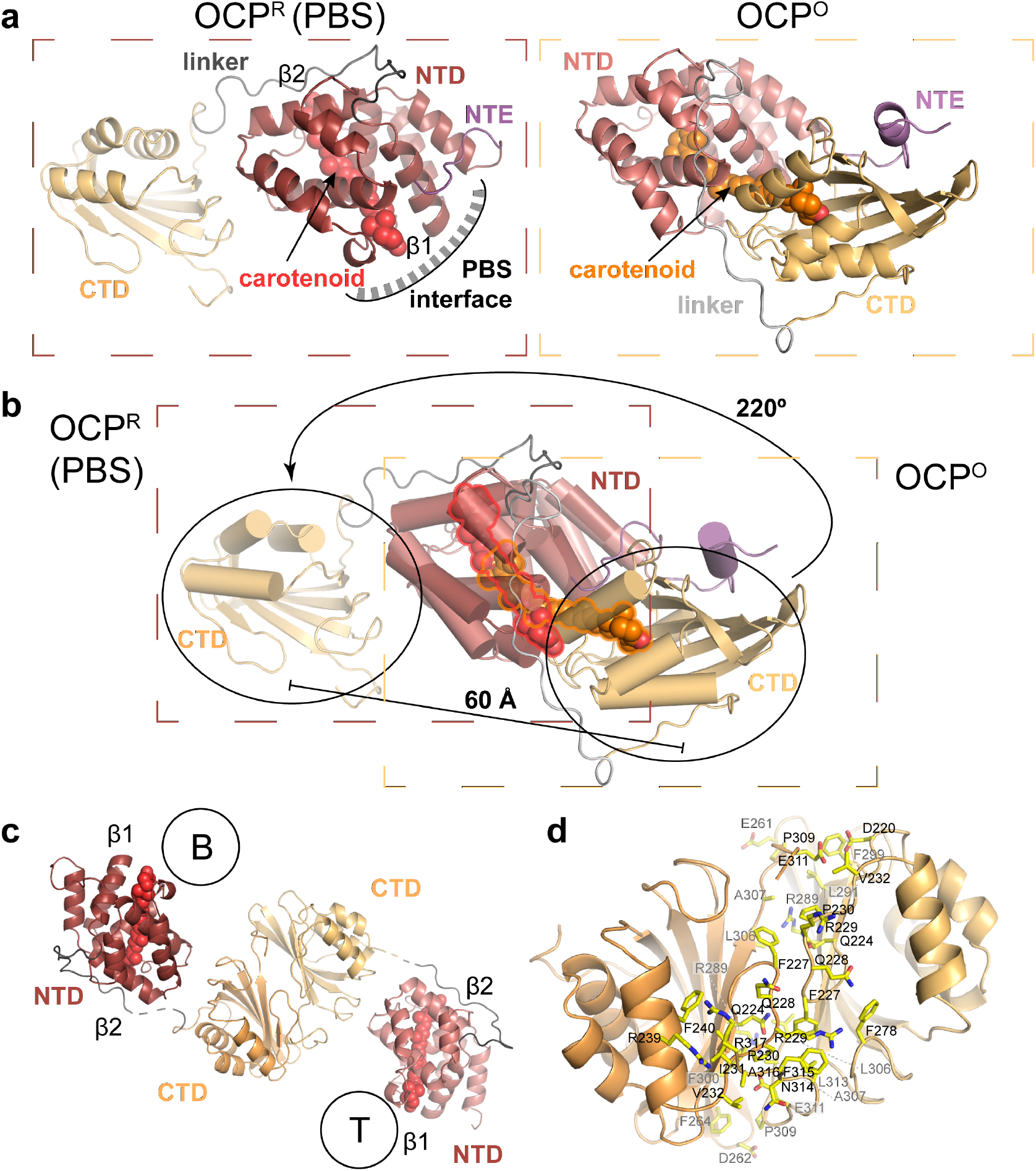
Structural changes associated with the photoconversion from OCP° to OCP^R^. **a**, Structure of OCP^R^ (left) in comparison to OCP° (right, PDB: 4XB5)^9^. β1 and β2 denote terminal rings of the carotenoid. **b**, Superimposed ribbon structures of OCP^R^ and OCP° in cartoon representation with helices as cylinders. CAN is shown as outlined spheres and colored in red for the OCP^R^ and orange for OCP°. The displacement of the CTD is indicated with an arrow. **c**, Structure of the OCP^R^ dimer via interaction of the CTDs. Circles with B and T represents bottom and top PBS cylinder, respectively. **d**, Close-up view of the CTD-CTD interface. Residues involved in the interaction within 4 Å are shown as yellow sticks. NTD = N-terminal domain, CTD = C-terminal domain, NTE = N-terminal extension.

We previously determined the X-ray crystal structure of the isolated NTD of OCP (PDB 4XB4)^9^, known as the Red Carotenoid Protein (RCP), that also can quench the PBS^9^. While the NTD of the OCP^R^ and the isolated RCP superpose with an rmsd of 0.4 Å over 121 aligned Cα atoms and exhibit a similar carotenoid conformation, the protein environment surrounding the chromophore differs due to the presence of the NTD-CTD linker (residues G166-T194) in OCP^R^. Residues E174-V177 are ordered across the β2 surface (Fig. 2 a) and are within 4 Å of the carotenoid. Moreover, the positioning of linker residue E174 possibly contributes to the pronounced asymmetry of the electrostatic distribution around the carotenoid. The net charge is negative around the β2 ring and positive around the β1 ring, which is proximal to a bilin (Extended Data Fig. 6), likely stabilizing the OCP^R^ in its binding site and fixing the mutual position of the carotenoid in OCP and the closest bilin in the PBS.

Unexpectedly, two OCPR, one bound to the B-cylinder, the other bound to the T-cylinder form a dimer interacting via their CTD domains (Fig. 2 c). The density for the CTD dimers is weak, indicating that they are mobile. The dimerization is in contrast quite stable, with a buried surface area of ∼1,150 Å^2^. The residues making up the dimerization interface are conserved in OCP (Fig. 2 d and Extended Data Fig. 7 a, c) and are shielded from solvent in the OCP° by interaction with the N-terminal extension (NTE, residues 1-18). Accordingly, the OCP^R^ structure connects two apparently disparate observations concerning structural changes associated with photoactivation: that OCP^R^ is a dimer in solution^15^, and that dimerization is mediated by residues exposed by dissociation of the N-Terminal extension during photoactivation^10,14^.

Reactivation of the quenched PBS depends on the 14 kDa fluorescence recovery protein (FRP), which interacts with the OCP-CTD^16,17^. Given the OCP^R^ dimers we observe in the quenching complex, we propose that an FRP dimer binds to the CTD-dimer. Each CTD could then form a complex with an FRP monomer while disrupting the CTD dimer, with FRP serving to protect the newly exposed hydrophobic residues on the CTD from solvent. The detached CTDs are then able to approach the NTD again to allow for re-association and formation of OCP°. This proposed mechanism is consistent with (i) previous data from controlled proteolysis experiments that showed that the CTD is not essential for quenching and was proposed to play a regulatory role^10^ and (ii) recent biochemical data suggesting that in catalyzing the reversion from OCP^R^ to OCP°, the OCP^R^ and FRP binding occurs in a 2:2 ratio^18^. One of the main differences of the conformation of the CTD in OCP^R^ is the loop formed by residues P276-W277-F278 which covers the entrance to the tunnel in the CTD that is occupied by carotenoid in OCP° (Extended Data Fig. 7 b). Based on an X-ray footprinting model^19^, the bound FRP could potentially also facilitate re-opening of the carotenoid tunnel for carotenoid re-entry by binding and repositioning those residues.

## The PBS-OCP interaction

Contrary to all previous predictions for the structure of the quenching complex, four OCP^R^ bind to the PBS core. The binding sites are located at the top and the bottom cylinders of the PBS (Fig. 1 a, b). Several, some mutually exclusive, models based on a wide range of experimental techniques have been proposed for the OCP-PBS interaction^20-26^. Some of the suggested gross characteristics of the interface, such as binding of the OCP to Allophycocyanin-A/B (ApcA/B, APC660)^22-24^ are confirmed and now precisely detailed by our structure (Fig. 1). OCP^R^ interacts mostly with the outermost ApcA/B disc in the T cylinder (Fig. 1 b, c). The two top binding sites (here named Ta and Tb) are identical other than minor differences in sidechain conformations. On the B cylinders, OCP^R^ is bound to the two ApcA/B discs with only a minor interaction with an ApcB from the ApcABEF disc (Fig. 1 b, c). Furthermore, our structure shows that OCP^R^ interacts with the CpcG1 rod-linker, a conserved subunit assumed to be present in the ancestor of the Cyanobacteria^27^, contributing to the surface of interaction (Fig. 1 c, 3 b Extended Data Table 2 and 3).

The interaction surfaces between the OCP^R^ and the B cylinders (∼1400 Å^2^) are identical and are slightly larger than the buried surface of interaction with the T cylinder (∼1200 Å^2^). The larger interaction surface of the B cylinder binding site results from additional interactions with the N-terminal extension (G10, I11, F12 and N14) and with the interdomain linker residues A165 and G166 with the ApcB (G21 and D25) from the ApcABEF disc (Fig. 3 a, b and Extended Data Table 2). A subset of the N-terminal extension residues released from the CTD upon conversion to OCP^R^ (Fig. 2a) interact with the B cylinder. Particularly, the conserved residue F12 shows extensive interactions with residues from the ApcA (highly conserved E76 and T80), ApcB (highly conserved Y62), and with P215 and M216 from the rod-core linker CpcG1 (Extended Data Table 2). The release of the N-terminal extension from the CTD in the conversion from OCP° to OCP^R^ not only enables the CTD-CTD dimerization, but also provides additional contact points to the PBS.

**Figure 3:**
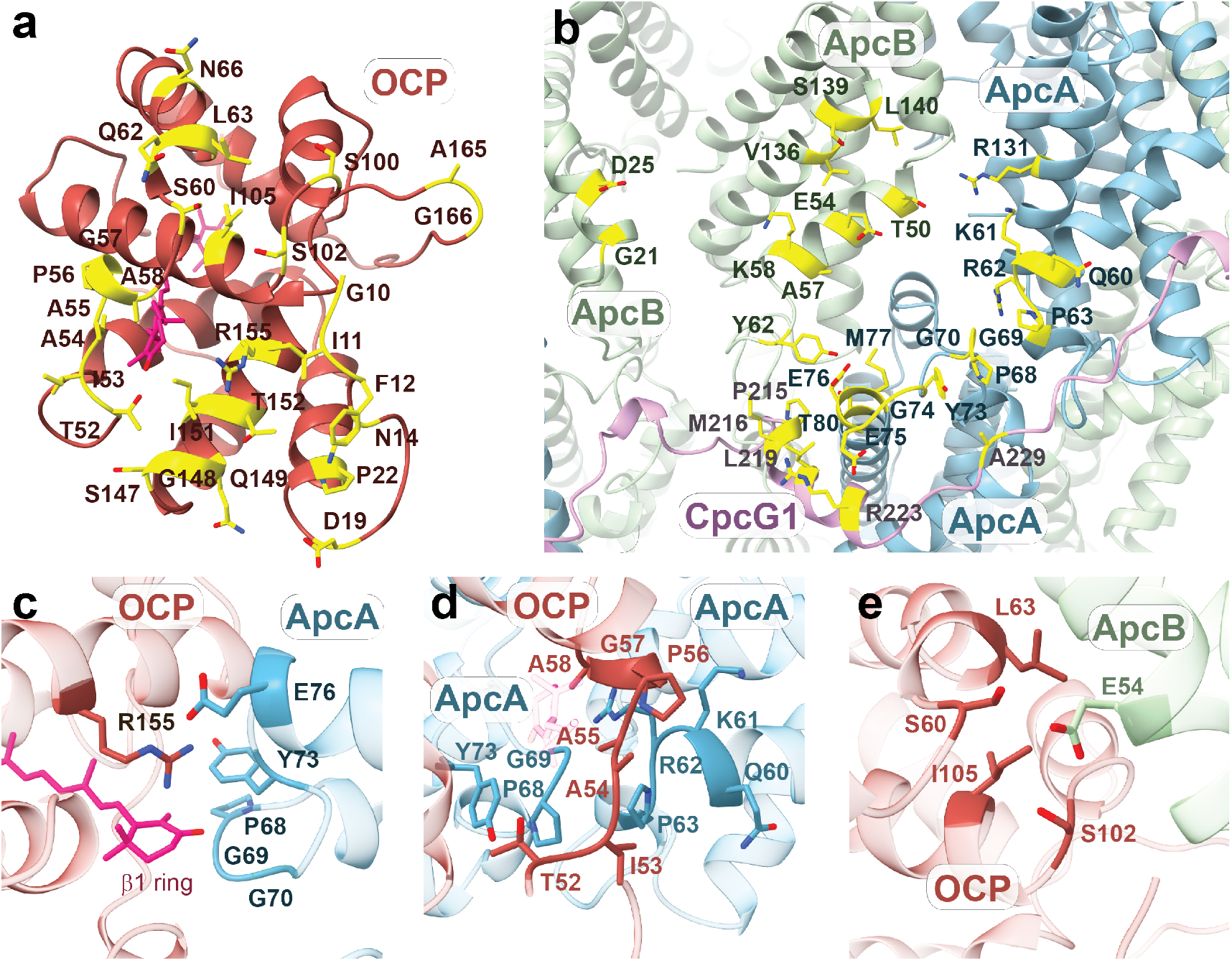
PBS-OCP interactions. **a and b**, Interface residues of OCP (a) and ApcAB and CpcG1 (b) within 4 Å of each other in the major interface shown in bottom cylinder. **c**, Close-up view of the interaction between the OCP surface surrounding the β1 ring of the carotenoid and ApcA (B cylinder). **d**, Close-up view of the interaction between the OCP loop (T52-A58) and ApcA (B cylinder). **e**, Interaction between E54 from ApcB and the OCP residues.

The majority of OCP^R^ residues involved in the interaction with the PBS are mainly from the β1 face of the NTD (Fig. 3 a, Extended Data Tables 2, 3). Residues from the CTD (N314, F315 and R317) are also in proximity to the PBS core but their density is very weak and therefore it is unlikely that those are specific interactions. The close vicinity does however explain why several studies using crosslinkers identified residues in the linker or in the CTD as interacting with PBS^20,21^. The conserved R155 from the NTD is in the center of the interaction surface next to the carotenoid and interacts with the highly conserved ApcA residue Y73 (Fig. 3 c). Additional interactions occur between R155 and E76 (conserved ApcA) at the B cylinder and G74, E76 and M77 (conserved ApcA residues) at the T cylinder (Extended Data Tables 2, 3). While most of the central interactions of OCP with PBS involve ApcA, there are also significant peripheral interactions with ApcB such as E54 that interacts with several residues from OCP (Fig. 3 e).

A further completely unexpected and large portion (∼300 Å^2^) of the OCP^R^ interface with the PBS is formed by the loop that connects the two discontinuous four-helix bundles that constitute the NTD. These residues, T52-A58, are predominantly small side chains that are strongly conserved (Fig. 3 d). The backbone of the OCP in this region closely approaches the ApcA backbone (G69) and between OCP residues (A55 and A58) with distances of only 5-6 Å explaining the conservation of small sidechains. Furthermore, the β1 ring of the CAN is surrounded by four highly conserved residues (P68, G69, G70 and Y73) from the ApcA (Fig. 3 c and Extended Data Tables 2, 3), stabilizing the center-to-center distance between both chromophores at ∼27 Å (Fig. 4 a, b).

**Figure 4.**
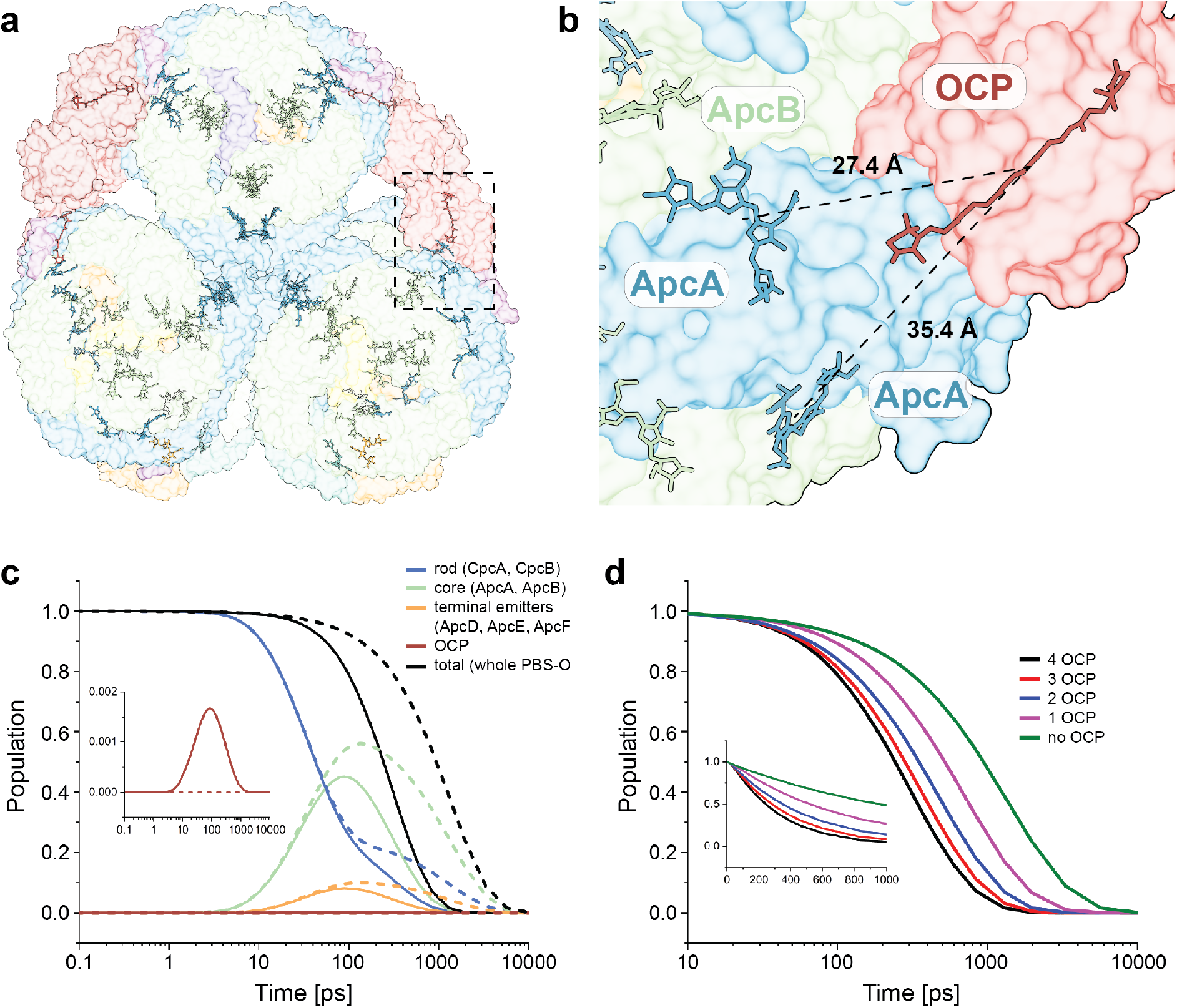
OCP-induced PBS quenching mechanism. **a**, Overview of the bilins and carotenoid (CAN). Dashed lines outline the region shown in panel b. **b**, Close-up view of the two closest carotenoid-bilin pairs with indicated distance. **c**, Simulated excitation energy flow in quenched (solid) and unquenched (dashed) OCP-PBS. The initial population is placed at the end of rod. The curves monitor the evolution of an excited state population of pigments located in rod, core, and terminal emitters. The black traces show the total population of the whole OCP-PBS complex. Inset shows the time profile of population of excited CAN in OCP, which is negligible due to the short (3.5 ps) CAN lifetime that is significantly shorter than the time constant of bilin-to-CAN energy transfer. **d**, Effect of the number of active OCPs on the total excitation lifetime in PBS. Inset shows the first nanosecond in linear time scale. Identical excitation conditions were used in all five simulation runs.

## Mechanism of the OCP-induced PBS quenching

We used the OCP-PBS structure to model the PBS quenching by the carotenoid bound to OCP^R^. We assume that only energy transfer is involved in the quenching mechanism^28^. Other potential mechanisms, such as excitonic quenching or electron transfer^26^, are unlikely given the ∼27 Å distance between the CAN in OCP^R^ and the closest bilin (Fig. 4 a, b). The simulation was carried out for the whole OCP-PBS complex, and the unquenched state of PBS was simulated by setting the transition dipole moment of CAN in OCP to zero, effectively switching off the OCP quenching (details of the model are provided in the Methods section).

The results of modeling the energy flow through the OCP-PBS complex in quenched and unquenched states are shown in Fig. 4 c. After excitation of a PBS rod, the population is transferred to the APC core within the first 100 ps and this process is not affected by OCP. The lifetime of the APC core is significantly shortened in the quenched state, proving that structural arrangement of OCP allows for efficient quenching of the core via energy transfer mechanism. The results show that for each of the four OCP CAN molecules there is one bilin characterized by bilin-to-carotenoid transfer time constant of 29±3 ps and one bilin transferring at 89±3 ps (for a CAN dipole moment of 2.3 D, see Extended Data Fig. 5 a, b). No other pairwise time constants shorter than 200 ps were found. These individual bilin-to-CAN quenching rates are much longer than the sub-ps value proposed by Tian et al.^29^. However, the values obtained from our model based on the presented OCP-PBS structure are clearly short enough to ensure efficient quenching that agrees with experimental data, but only under conditions where four OCPs are bound to PBS. If we leave only one, two or three OCPs active, the quenching efficiency decreases significantly (Fig. 4 d, Extended Data Fig. 5 c, d). Thus, the key structural feature ensuring efficient quenching via energy transfer mechanism is the binding of two dimers of the OCP^R^ to the PBS. Moreover, our modeling suggests that OCP bound to one PBS within an array can quench energy captured by a different array PBS due lateral energy transfer, making the array arrangement efficient at low light and at same time rapidly responsive when exposed to higher light intensities (Extended Data Fig. 5 e, f).

The efficient quenching is important for fitness of the organism in its natural environment, but the overall NPQ process is counterproductive in cyanobacterial mass cultures, decreasing the photosynthetic sunlight-to-biomass conversion efficiency by up to 60% ^2,30^. Indeed, tuning the balance between light harvesting and photoprotection is arguably the greatest challenge in realizing the full potential of cyanobacteria as solar powered cellular factories^31^. The structure of the quenched state of a cyanobacterial PBS provides the requisite description of excitation energy transfer in the quenching process to inform efforts to modify the PBS, OCP and/or FRP to alleviate this energetic wasteful process^30,32^ while at the same time providing the first high resolution model of a quenched light harvesting antenna with implications for understanding and engineering both natural and articial photosynthetic systems.

## Methods

### Preparation of the *Synechocystis* PCC 6803 Phycobilisome

*Synechocystis* PCC 6803 were grown photoautotrophically in a BG11 medium. Cells were kept in a rotary shaker (100 rpm) at 30°C, under 3% CO_2_ enrichment, illuminated by white fluorescent lamps with a total intensity of about 30 μmol photons m^−2^ s^−1^. The protocol used for PBS isolation was based on ^33,34^ and the procedure as described in ^11^.

### Expression and purification of *Synechocystis* PCC 6803 OCP-CAN holoprotein in *E. coli*

A C-terminal 10x His-tagged *Synechocystis* PCC 6803 *ocp* gene (*slr1963*) was cloned in a pCDFDuet (Novagen), resulting in a plasmid named pEP4. Then, BL21(DE3) cells were transformed simultaneously with pEP4 and pAC-CANTHipi plasmid (Addgene plasmid #53301) and induced with 100 μM of Isopropyl ß-d-1-thiogalactopyranoside (IPTG) and incubate overnight at 25 °C. OCP-CAN was firstly purified by a Ni-NTA affinity chromatography (HisTrap affinity column, GE Healthcare), yielding a mixture of apo-and holo-OCP protein. Then, to separate the holoprotein from the apoprotein, a hydrophobic interaction chromatography using a linear decreasing (NH_4_)_2_SO_4_ gradient (from 1.5 M to 0 M) (HiTrap HIC phenyl column, GE Healthcare) was applied to the mixture. The maximum extinction coefficient of CAN (118,000 M^−1^cm^−1^; 1:1 binding stoichiometry of CAN:OCP) was used to determine the concentration of the OCP-CAN holoprotein. The absorption spectrum of the *E. coli-*expressed 6803 OCP inactive and photoconverted to the red form is consistent with previous literature report^9^ (Extended Data Fig. 4).

### Protein separation

The isolated PBS samples were treated as described in ^11^. The purity of isolated OCP protein was assessed by Coomassie Brilliant Blue stained SDS-PAGE (Extended Data Fig. 4).

### Absorption and fluorescence spectrum measurement

OCP-CAN holoproteins were buffer-exchanged into 50 mM Tris-HCl, pH 8.0, and 200 mM NaCl before spectroscopic measurements. PBS and OCP-CAN absorption spectra were collected with a Cary 60 spectrophotometer (Agilent). Photoconversion of the inactive OCP° samples to the active OCP^R^ form was carried out with 15 min of blue LED illumination (λmax= 470 nm, Philips Lumileds LXML-PB01-0030) at 1,000 μmol photons m^−2^ s^−1^. The fluorescence emission spectra of the PBS were recorded at room temperature from 600 to 800 nm in a fluorimeter (TECAN Spark 20M multimode microplate reader) with an excitation wavelength of 580 nm.

### In vitro OCP-PBS biotinylated complex formation

The PBS sample was biotinylated as described in ^11^. The OCP was previously illuminated for 10 minutes by blue-LED light (λmax= 470 nm, 1,000 μmol photons m^−2^ s^−1^) to activate it. Then, for OCP-PBS complex formation, activated OCP was added to the isolated biotinylated PBS (in 0.75 M potassium phosphate pH 7.5), giving an OCP to PBS ratio of 20. The samples were illuminated again under the same light for 20 minutes at room temperature. Fluorescence spectra were measured to quantify the OCP-induced quenching (Extended Data Fig. 4).

### Cryo-EM sample preparation of OCP-PBS

Cryo-EM sample preparation was carried out as described in^11^. For sample preparation, we used Quantifoil Au 300 mesh 2/1 grids covered with a home-made streptavidin monolayer, which were manufactured as described previously^35^. To apply the sample, grids were first rehydrated in buffer A (375 mM potassium phosphate, pH 7.5) and then blotted dry with filter paper. To promote the conversion of OCP° to OCP^R^ the OCP-PBS sample was illuminated with blue light at a wavelength of 470 nm for 5 min on the bench prior to application to the grid. 4 μl of OCP^R^-PBS at a concentration of 5.3 mg/ml (0.86 μM) in buffer B (750 mM potassium phosphate, pH 7.5) were added to the grid and then incubated on the bench for 60 seconds. Grids were washed on two 10 μl drops of buffer C (375mM potassium phosphate, pH 7.5, 3% w/v trehalose, 0.01% v/v NP40, 0.05% w/v beta-octylglucoside) before being carefully wicked with Whatman filter paper. 1μl of buffer C was added immediately, and the grid lifted into a FEI Mark IV Vitrobot. The grid was then manually blotted for 2-3 s at 18°C and 100% humidity before plunging into a liquid ethane-propane (3:1) mix^36^. During the process the OCP-PBS was exposed to ambient light for a total of approximately 2-3 minutes between pipetting and vitrification, in addition to the initial illumination.

### Cryo-EM data collection OCP-PBS

The data collection pipeline used in this study was the same as for the PBS without OCP^11^. 13427 movies were collected on a Titan Krios G3i microscope operating at a voltage of 300 kV, equipped with a Gatan K3 direct electron detector operating in CDS mode^37^ and a GIF Quantum energy filter with a 20-eV slit width. Movies were acquired with SerialEM^38^ in super-resolution counting mode with a super-resolution pixel size of 0.525 Å using an image shift collection scheme with active beam tilt correction, a defocus range from -0.5 μm to -1.6 μm and a total exposure of 50 e^-^/Å^2^.

### Image processing OCP^R^-PBS

The image processing pipeline was similar to PBS without OCP^11^. All super-resolution movies were aligned, gain corrected and binned by 2 using MotionCorr2 as implemented in RELION3^39^. The background streptavidin lattice in the motion corrected micrographs was subtracted using in-house scripts^35^. These subtracted micrographs were then imported into CryoSPARC^40^ for patch CTF estimation and further image processing.

Particles were picked from a subset of 870 micrographs using templates generated from the PBS^up-up^ reconstructions from dataset 1 of the PBS without OCP^11^ and with a box size of 1000 pixels. Heterogeneous refinement yielded a single good class that reached 3.4 Å resolution during refinement. Subsequently, particles were picked from all micrographs using the same templates and a box size of 720 pixels. From a total of ∼2,500,000 particles ∼1,100,000 particles classified into a single good class which corresponded to PBS-OCP^R^, with no other conformations in the dataset apparent. Inspection of the reconstruction revealed flexibility in the core region, resulting in elongated density artifacts in certain parts of the map and smeary density for OCP. To speed up subsequent processing steps the boxsize was cropped to 360 pixels in real space, thus excluding the peripheral rods. 3D variability analysis (3DVA) revealed an overall twisting motion of the core and the partial loss of the bottom ApcAB hexamer rings on either side, presumably because of the overall fragility of the complex. A subset of particles showing improved OCP density was identified using 3DVA in cluster mode. This subset (∼341,000 particles) was subjected to non-uniform refinement to yield a final resolution of 2.6 Å.

This particle stack was used to further improve several features of the core. To further improve the density for OCP, particle subtraction followed by local refinement was performed to obtain a map of OCP bound to the top ApcAB double hexamer at 2.6 Å resolution with the map quality for OCP visibly improved. To improve the C-terminal domain of OCP, this latter map was analyzed by 3DVA and a particle subset with better density for CTD, but a slightly worse overall resolution was identified (∼180,000 particles, 2.8 Å resolution).

Similar strategies were used to obtain locally refined maps of the entire top cylinder (2.6 Å) and the entire B2 cylinder (2.5 Å), each with improved map quality at the same nominal resolution. To obtain a reconstruction of the holo OCP^R^-PBS including the rods the cropped particles were re-extracted with a box size of 720 pixels and subjected to homogenous refinement, yielding an overall resolution of 2.7 Å.

To obtain a high-resolution reconstruction of the rods, templates obtained from dataset 1 from the PBS sample without OCP were used to pick and extract particles with a box size of 360 pixels. After 2D classification a set of ∼2,100,000 particles was selected for further processing. CTF refinement and non-homogenous refinement yielded a reconstruction with a resolution of 2.1 Å, thus reaching Nyquist limit. Further classification of the particle set to probe the possible presence of different rod protomers within one PBS complex always yielded the same conformation.

All processing was carried out using C1 symmetry and resolutions were estimated with the FSC 0.143 criterion. For interpretability and illustrations, reconstructions were sharpened using DeepEMhancer ^41^ with the default tight mask preset.

### Atomic model building and refinement PBS-OCP^R^

Atomic models were built by first fitting existing structures of the α and β-PC subunits into the density for orientation (4F0T and homology models based on algal homologs from 5Y6P/6KGX) that were then fit into the density using COOT 0.95 (ref. ^42^). Linker domains were built ab initio by tracing the backbone and assigning the sequence and fitting the sidechains using COOT 0.95 (ref. ^42^). Models were built sequentially, starting with the maps of the locally refined B- and T-cylinders and the locally refined T-disc 2. The resulting atomic models were refined using the map of the core with appropriate restraints with the real space refinement program in PHENIX 1.19.2 (ref. ^43^). The rods were modelled and refined independently using the same approach. To arrive at a model for the entire PBS-OCP^R^ complex, maps and models for the individual rods were rigid-body docked into the density of the holo-PBS-OCP^R^ complex, which resulted in an unambiguous orientation of each rod. Because the holo-PBS-OCP^R^ maps are not resolved enough in the distal rod regions to allow for model refinement, models of the full complex are for visualization only.

### Model of PBS Quenching

The model used to calculate energy transfer in OCP-PBS complex is the same as described in our accompanying study^11^. Here, the key modification is the inclusion of the CAN bound to OCP. For the transition dipole moment of the S_0_-S_1_ transition of CAN (because the S_1_ state should be the energy acceptor), we used the value of 2.3 D obtained from calculations using combination of molecular modelling and quantum chemistry^44^. We note that this is a mean value obtained for CAN in OCP°. Thus, the actual value for OCP^R^ bound to PBS may differ, but we note that binding to OCP increased the mean dipole from 1.7 D in THF solution to 2.3 D in OCP^44^. The significant asymmetry in charge distribution around CAN when OCP is bound to PBS (Extended Data Fig. 6) is comparable to that in OCP°,^45^ justifying our approximation. The effect of the actual value of the carotenoid transition dipole moment were also studied, as shown in Extended Data Fig. 5 a, b. As expected, larger S_1_ dipole moment increases the efficiency of excitation quenching. To calculate the spectral overlap, the hypothetical absorption spectrum of the acceptor, which is associated with the forbidden S_0_-S_1_ transition of CAN, was modelled as a mirror image of the S_1_ emission spectrum of the carotenoid peridinin reported earlier^46^. The resulting spectrum was then shifted to match the expected 0-0 energy of carotenoid in OCP^47^. The CAN S_1_ lifetime of 3.5 ps was used in simulation, based on the value reported in Kuznetsova et al.^45^. However, since the S_1_ lifetime of CAN is much shorter than the bilin-to-carotenoid quenching time, the actual value of the S_1_ lifetime has little effect on quenching dynamics. To evaluate the validity of the quenching model, we compared simulated decay-associated spectra (see ^11^ for details) with those obtained from fitting the experimental data^29^. The results are shown in Extended Data Fig. 5 c, d, demonstrating reasonable match between our model based on energy transfer quenching and experimental data obtained from time-resolved fluorescence.

### Bioinformatics

We obtained HMM profiles for each phycobiliprotein by collecting all phycobilisome sequences using jackhmmer (https://www.ebi.ac.uk/Tools/hmmer/search/jackhmmer) against the rp75 Uniprot proteome with a starting sequence of each *Synechocystis* PCC 6803 phycobilisome protein. Apc and linker proteins were then each aligned using ClustalW^48^ and used to generate a phylogenetic tree using RAxML^49^ with default parameters and PROTGAMMAW substitution matrix. Sequences from major branches that contain each *Synechocystis* PCC 6803 protein were aligned with ClustalW and trimmed with trimAl^50^ and the trimmed alignment was used to generate HMM profiles for each protein. The HMM profiles were then combined into a phycobilisome sequence HMM library. We then downloaded the 500 most complete non-redundant cyanobacterial proteomes from Uniprot and searched them with the combined HMM profile library and occurrences of each type of phycobilisome protein were counted. We then compared the protein counts with those of *Synechocystis* PCC 6803 and chose the 50 closest proteomes for sequence conservation analysis. For each phycobilisome protein we then aligned those sequences with ClustalW^48^, manually checked the alignment, trimmed them with trimAl^50^ and visualized the protein sequence conservation with Weblogo 3.74^51^.

### Structural visualization and accession codes

Structures were analyzed with Chimera^52^, ChimeraX^53^ and Pymol (The PyMOL Molecular Graphics System, version 1.7 Schrödinger, LLC). Protein interfaces were analyzed using PISA^54^ and Profunc^55^ at the European Bioinformatics Institute. The pymol APBS plugin^56^ was used with default parameters to calculate the electrostatics surfaces shown in Extended Data Fig. 6.

The atomic coordinates have been deposited in the Protein Data Bank with the accession codes 7SC9, 7SCB, 7SCC, 7SCA.

The EM maps have been deposited in the Electron Microscopy Data Bank with the accession codes EMDB-25030, EMDB-25031, EMDB-25032, EMDB-25033 and EMDB-25068.

All other data are available from the corresponding authors upon reasonable request.

## Acknowledgements

CAK and MADM dedicate this manuscript to the late Dr. Nicole Tandeau de Marsac. Molecular graphics images were produced using the UCSF Chimera package from the Resource for Biocomputing, Visualization, and Informatics at the University of California, San Francisco (supported by NIH P41 RR-01081). DB and TP thanks the Czech Science Foundation, grant No. 19-28323X. DB also acknowledges institutional support RVO:60077344. Research in the Kerfeld lab was supported by the Office of Science of the U.S. Department of Energy DE-FG02-91ER20021. This project has received funding from the European Union’s Horizon 2020 research and innovation programme under the Marie Sklodowska-Curie grant agreement No. 795070. EN is a Howard Hughes Medical Institute Investigator.

## Author contributions

MADM and PVS designed and performed experiments, interpreted results. CAK designed and supervised the project. HK helped with the sample preparation, interpreted results. MS refined the structures and interpreted results. DB and TP performed the model for the quenching. BJG performed initial characterization of the specimen for cryo-EM. MADM, PVS and CAK wrote the manuscript with help from all authors.

## EXTENDED DATA

**Extended Data Fig. 1:**
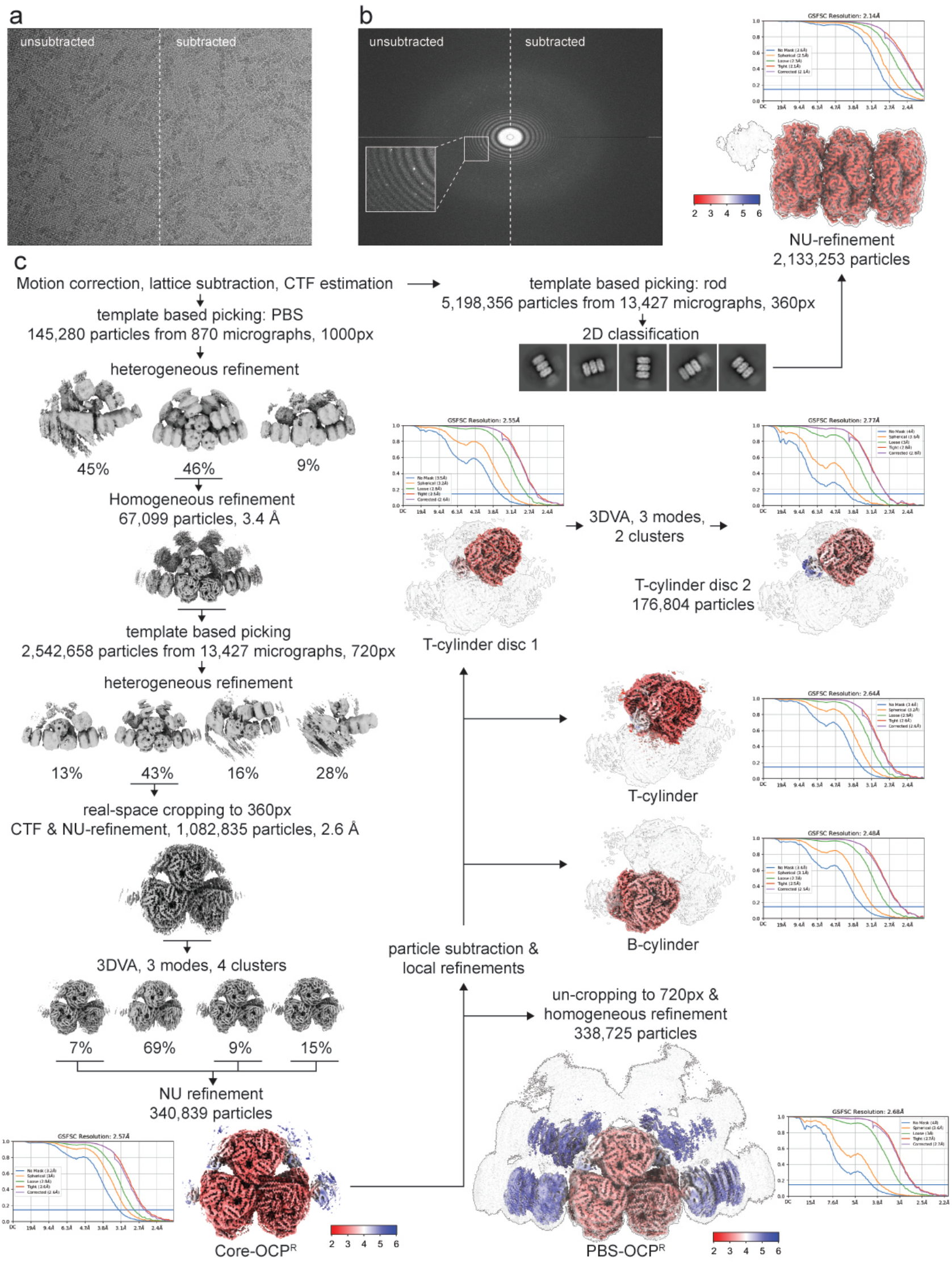
Cryo-EM analysis of the OCP-PBS complex. **a**, Example raw micrograph before and after streptavidin lattice subtraction. **b**, Fourier transform of (a) showing Bragg diffraction from the streptavidin crystal before subtraction. **c**, The workflow for the cryo-EM data processing. Red to blue color shading represents local resolution estimates at FSC=0.5 ranging from 2 to 6 Å. Maps of PBS-OCP and the rod are shown with two different thresholds to show flexible regions and connectivity.

**Extended Data Fig. 2:**
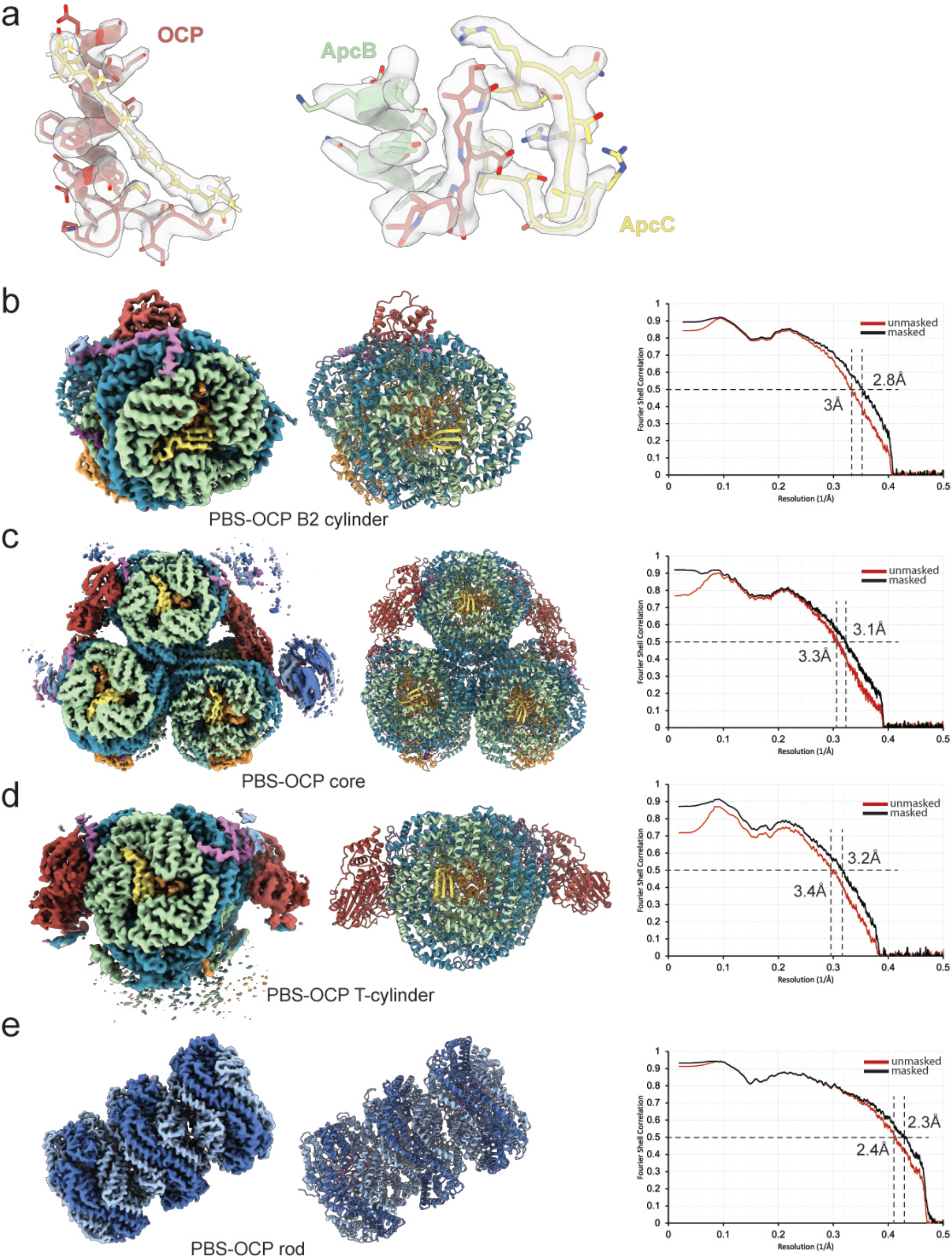
Map and Model quality of PBS-OCP^R^. a, examples of map density quality from different regions of the PBS-OCP^R^. b-e, Map, model and corresponding Fourier shell correlation (FSC) curves. Resolution of masked and unmasked models are indicated at FSC = 0.5.

**Extended Data Fig. 3.**
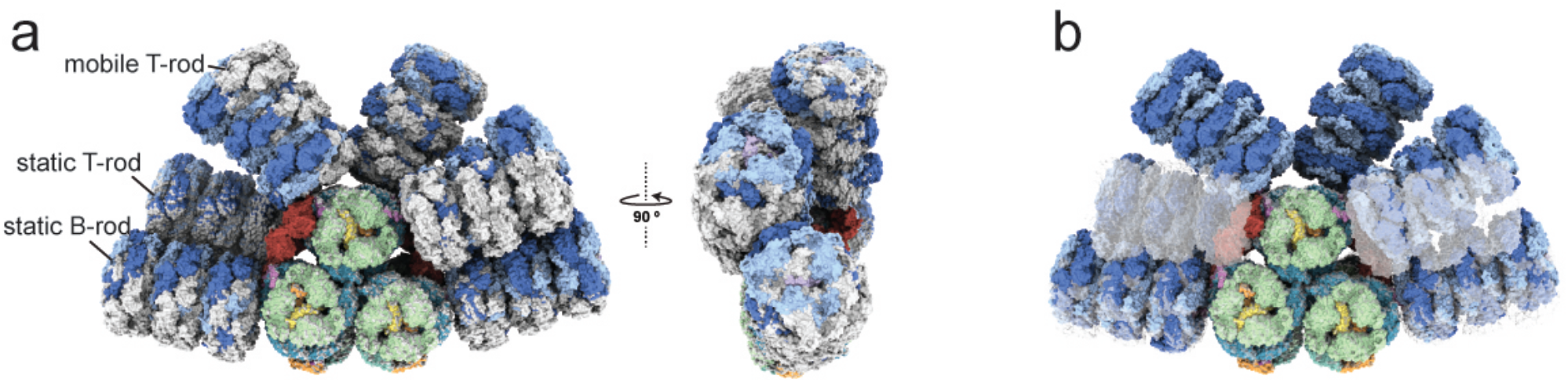
Structural comparison between light-harvesting PBS and quenched PBS. **a**, Superimposition of the PBS-OCP^R^ (colored as in Fig. 1) and PBS up-up (grey). **b**, Superimposition of the PBS-OCP^R^ and the PBS down-down conformation (transparent surface). Both shown in surface representation.

**Extended Data Fig. 4:**
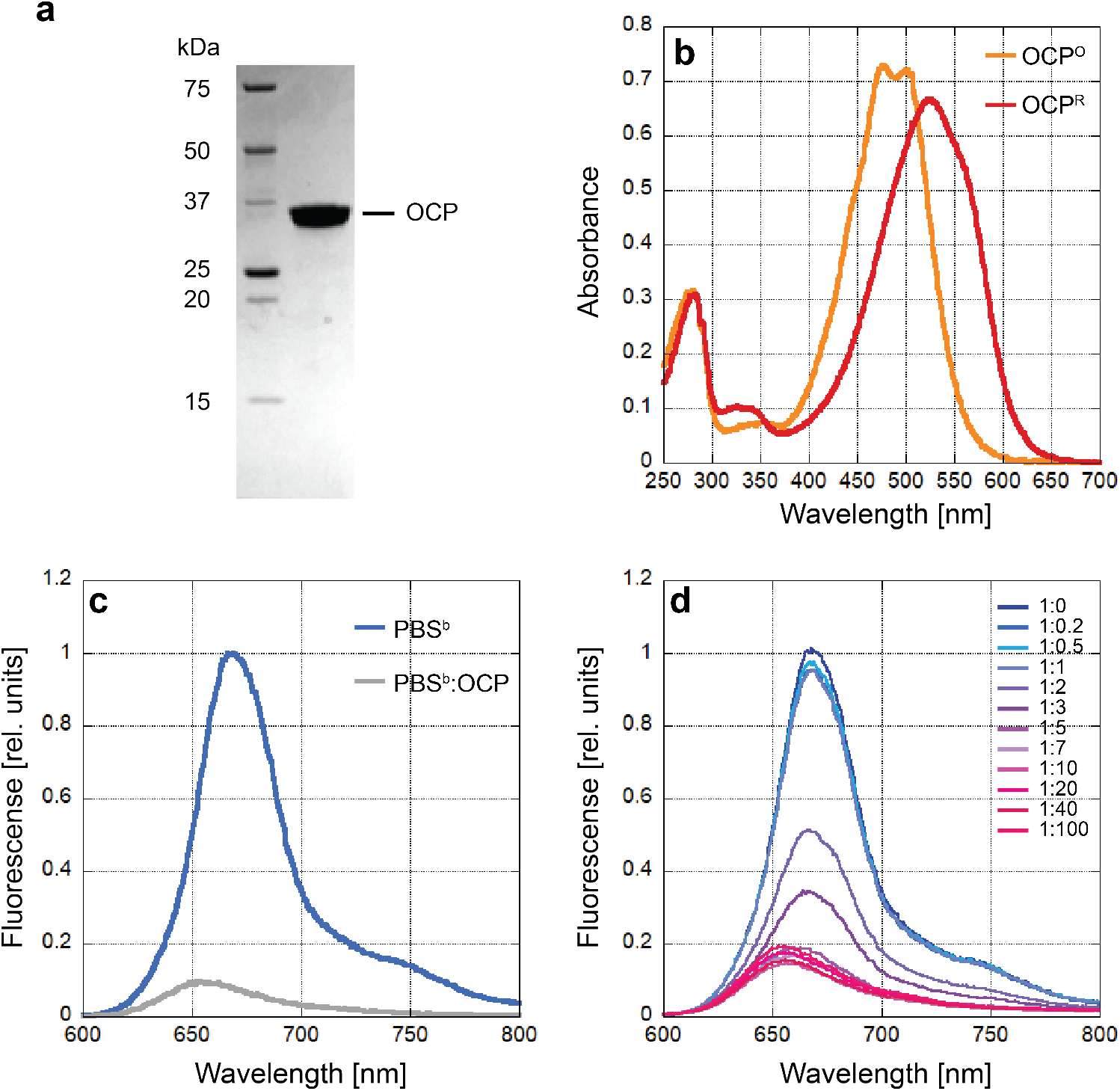
*Synechocystis* PCC 6803 OCP Purification. **a**, Coomassie blue-stained SDS-PAGE. **b**, UV-Vis absorption spectra of the OCP° (inactive form) and OCP^R^ (active form). **c**, Fluorescence quenching of 1:20 (PBS:OCP). The PBS is biotinylated. **d**, Titration PBS:OCP fluorescence quenching at various ratios indicated.

**Extended Data Fig. 5.**
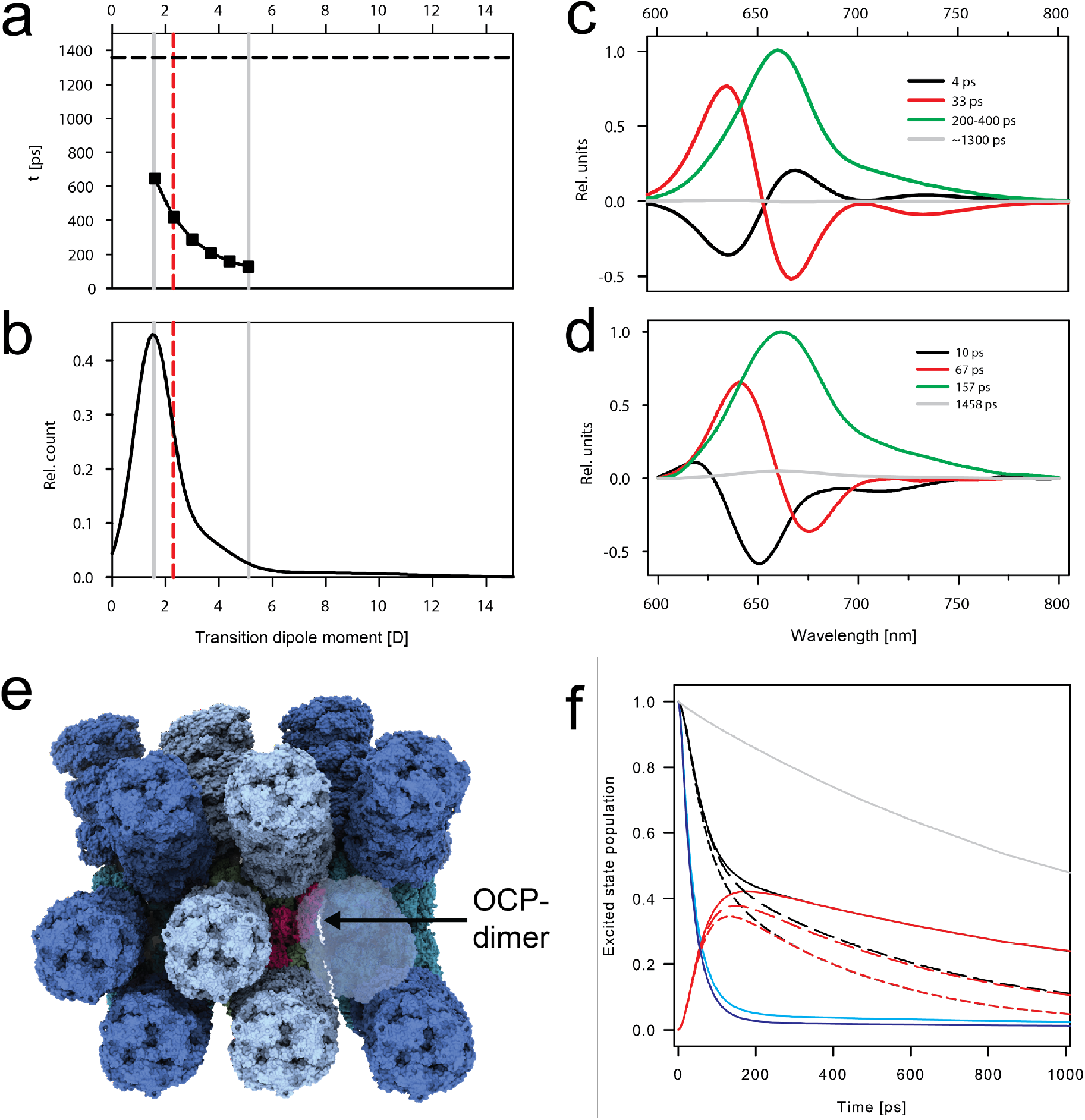
**a**, Effect of the transition dipole moment (TDM) of canthaxanthin in OCP on the lifetime of excitation in PBS; the dashed line indicates the lifetime of unquenched PBS (TDM = 0); **b**, Distribution of TDM values of echinenone in OCP, adapted from ^44^. The red line indicates the mean value, 2.3 D; the range marked by grey lines is used for simulations shown in panel (a). **c**, Decay associated spectra obtained from a simulation based on the OCP-PBS structure. It represents the excitation energy flow in the fully quenched OCP-PBS after excitation into the far end of the rod. The value of the green component, corresponding to the quenching, depends on the TDM value shown in panel (a). **d**, The same spectra obtained from fitting the time-resolved fluorescence data (adapted from^29^). **e**, PBS array with OCP bound. OCP biding sites are still accessible. **f**, Simulation of excitation energy flow in a phycobilisome pair (PBS1, black; PBS2, red) after excitation into the outer tip of the PBS1 bottom rod. Cyan (single PBS) and blue (PBS pair) kinetics monitors population in the rod. Grey, black and red lines represent excitation summed over all bilins in the PBS. Grey line shows the total excitation in a single PBS without OCP; In the PBS pair, the excitation is distributed from the initially excited PBS1 (black) to PBS2(red). Solid line, PBS pair without OCP; short dash, each PBS binds 4 OCPs; long-dash PBS2 binds 4 OCPs, PBS1 has no OCP. This simulation shows that the effect of OCP is shared between PBS in the pair. Analysis of transfer times suggested less than 20 inter-PBS bilin pairs with transfer times shorter than 20 ps. Majority of those were in the top cylinder of the core. Several close contacts were also identified between rods, however, the comparison of the blue and cyan traces shows that arranging PBS into a pair has little effect on the excitation dynamics within a rod.

**Extended Data Fig. 6:**
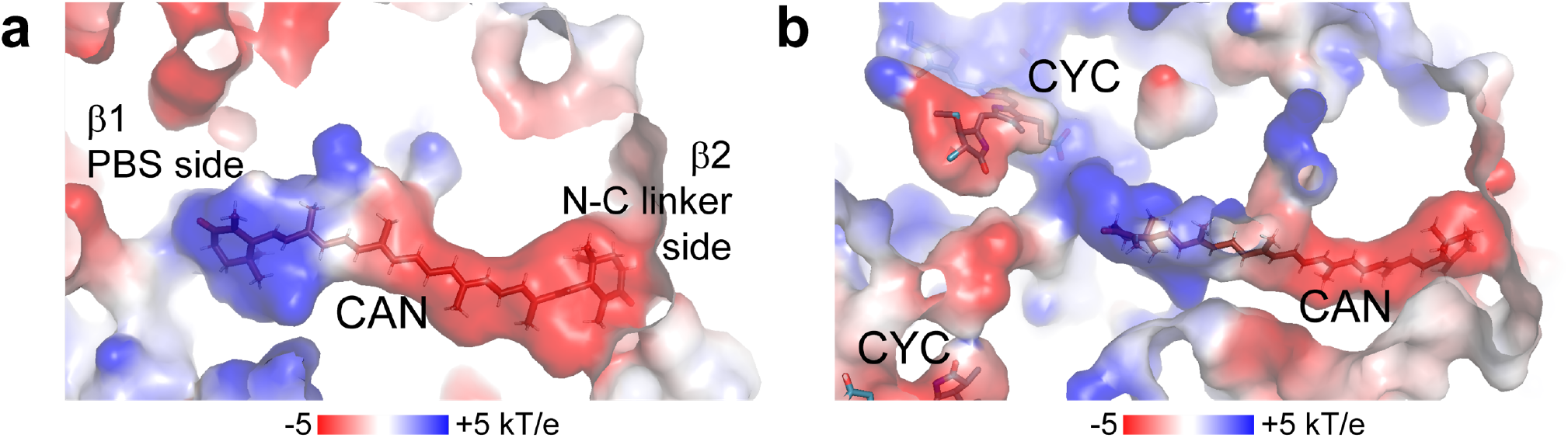
Electrostatics of the protein environment of the carotenoid tunnel mapped on the OCP^R^ bound to the PBS. **a**, Carotenoid tunnel with the β1 and β2 indicated and carotenoid shown as sticks. **b**, Electrostatic surfaces of the proteins around the carotenoid tunnel and the nearby bilins; carotenoid and bilins shown as sticks.

**Extended data Fig. 7:**
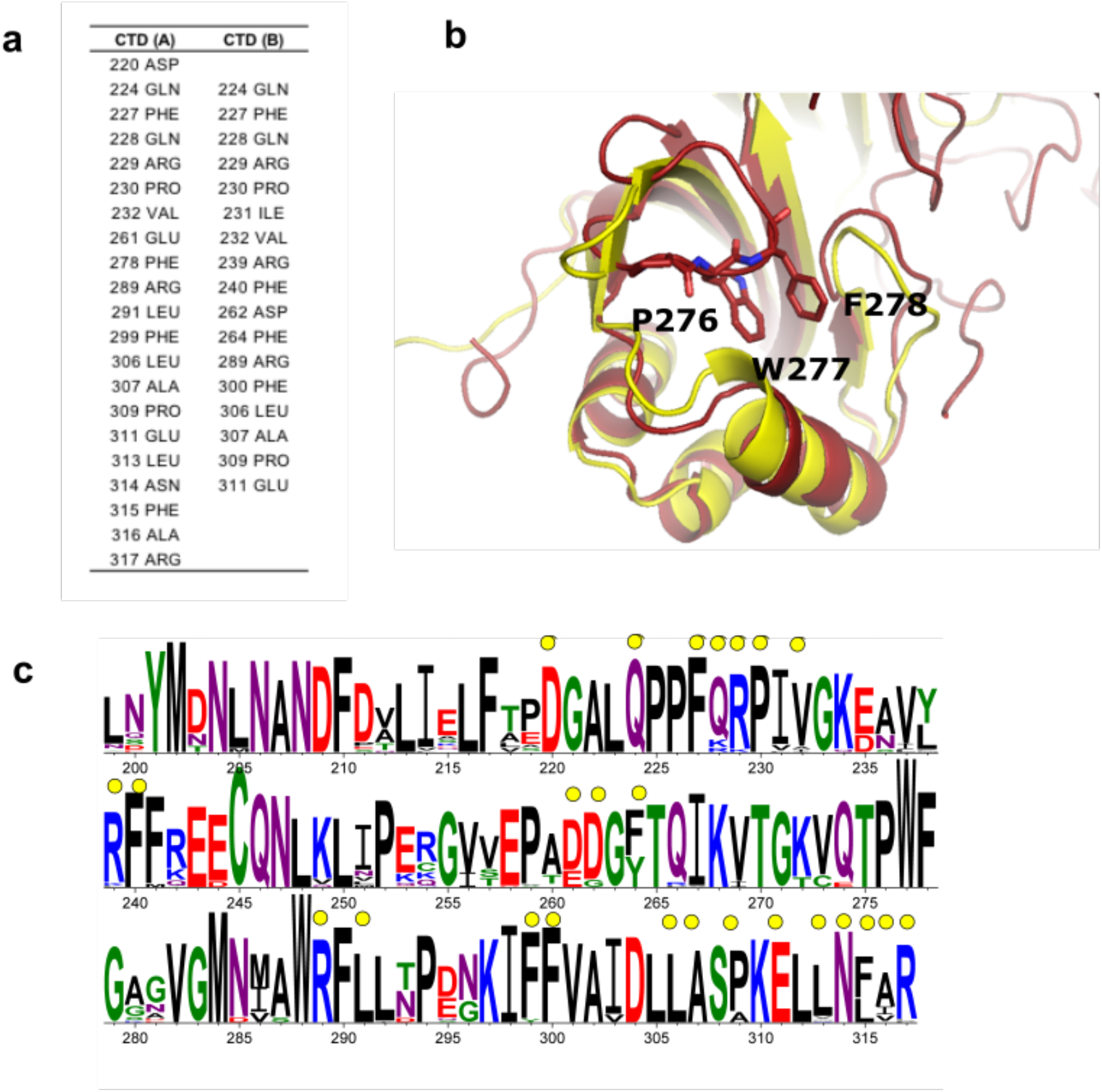
CTD from OCP. **a**, List of residues that interact within 4 Å. **b**, Close-up view of the P276-W277-F278 loop covering the opening to the carotenoid tunnel in the CTD of OCP°. **c**, Sequence conservation logo for the CTD (beginning at residue 199) of the OCP. Residues involved in the dimerization are highlighted with a yellow dot.

**Extended Data Table 1:**
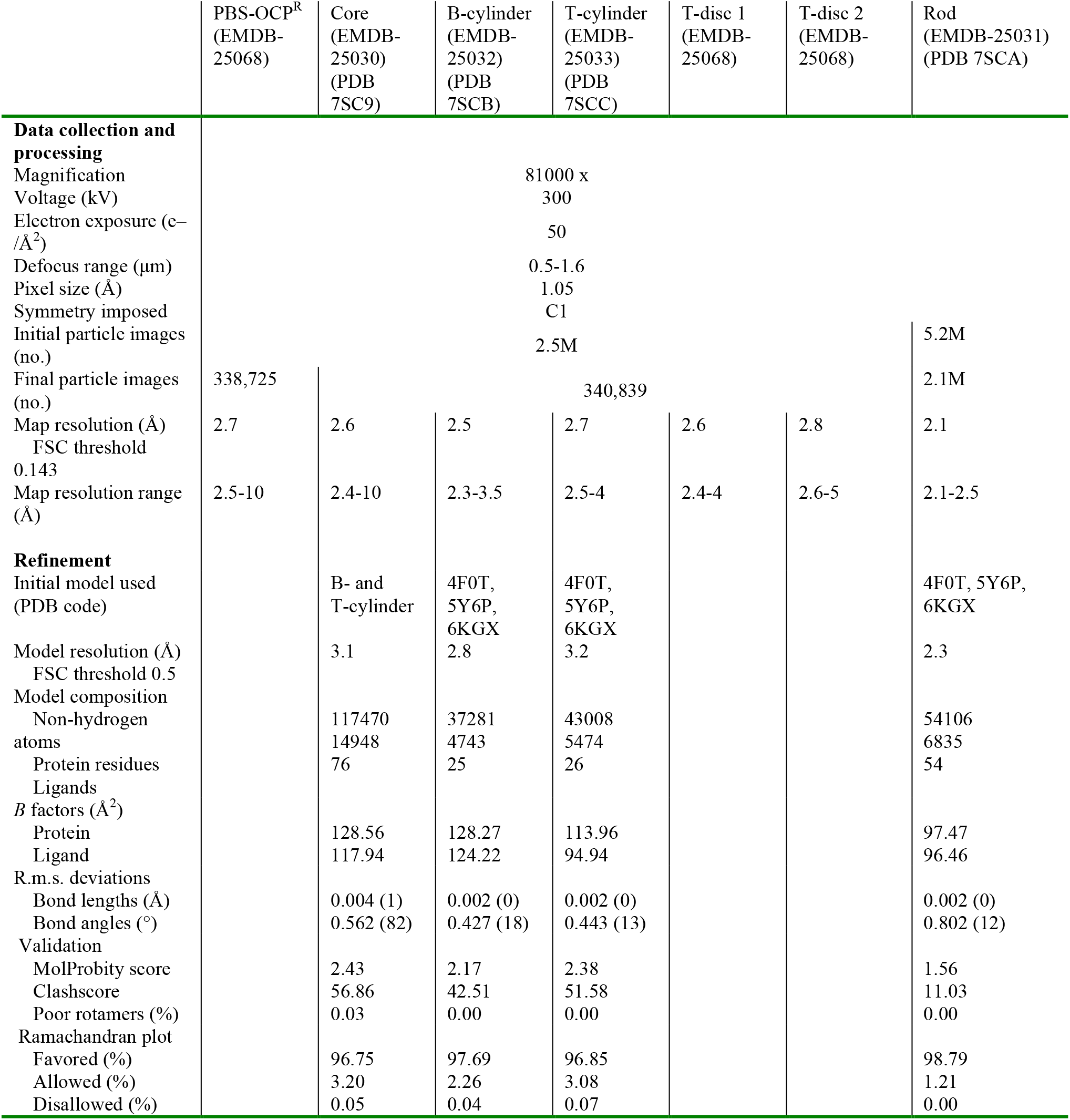
Cryo-EM data collection, refinement and validation statistics.

**Extended Data Table 2:** Pairwise interactions within 4 Å between the OCP and the B cylinders.

| **OCP** | **PBS** |  |  |
| --- | --- | --- | --- |
| *Residue* | *Residue* | *Trimer/protein* | *Chain* |
| Gly10 | Tyr62 | b23 | D (ApcB) |
|  | Ala57 | b23 | D (ApcB) |
| Ile11 | Glu76 | b23 | E (ApcA) |
|  | Met77 | b23 | E (ApcA) |
|  | Tyr62 | b23 | D (ApcB) |
| Phe12 | Tyr62 | b23 | D (ApcB) |
|  | Glu76 | b23 | E (ApcA) |
|  | Thr80 | b23 | E (ApcA) |
|  | Pro215 | cpcg1 |  |
|  | Met216 | cpcg1 |  |
| Asn14 | Leu219 | cpcg1 |  |
|  | Arg223 | cpcg1 |  |
| Asp19 | Arg223 | cpcg1 |  |
| Pro22 | Arg223 | cpcg1 |  |
| Thr52 | Tyr73 | b23 | E (ApcA) |
| Ile53 | Pro68 | b23 | E (ApcA) |
|  | Pro63 | b24 | E (ApcA) |
| Ala54 | Gln60 | b24 | E (ApcA) |
| Ala55 | Gly69 | b23 | E (ApcA) |
|  | Pro68 | b23 | E (ApcA) |
|  | Lys61 | b24 | E (ApcA) |
|  | Pro63 | b24 | E (ApcA) |
| Pro56 | Lys61 | b24 | E (ApcA) |
| Gly57 | Lys61 | b24 | E (ApcA) |
|  | Arg62 | b24 | E (ApcA) |
| Ala58 | Gly69 | b23 | E (ApcA) |
|  | Arg62 | b24 | E (ApcA) |
| Ser60 | Thr50 | b23 | D (ApcB) |
|  | Glu54 | b23 | D (ApcB) |
| Gln62 | Thr50 | b23 | D (ApcB) |
|  | Leu140 | b23 | D (ApcB) |
|  | Arg131 | b24 | E (ApcA) |
| Leu63 | Glu54 | b23 | D (ApcB) |
|  | Val136 | b23 | D (ApcB) |
|  | Ser139 | b23 | D (ApcB) |
|  | Leu140 | b23 | D (ApcB) |
| Asn66 | Ser139 | b23 | D (ApcB) |
| Ser100 | Lys58 | b23 | D (ApcB) |
| Ser102 | Glu54 | b23 | D (ApcB) |
| Ile105 | Glu54 | b23 | D (ApcB) |
| Ser147 | Tyr73 | b23 | E (ApcA) |
|  | Ala229 | cpcg1 |  |
| Gly148 | Tyr73 | b23 | E (ApcA) |
|  | Gly74 | b23 | E (ApcA) |
|  | Glu75 | b23 | E (ApcA) |
| Gln149 | Glu75 | b23 | E (ApcA) |
| Ile151 | Tyr73 | b23 | E (ApcA) |
|  | Gly74 | b23 | E (ApcA) |
| Thr152 | Gly74 | b23 | E (ApcA) |
|  | Glu75 | b23 | E (ApcA) |
|  | Glu76 | b23 | E (ApcA) |
| Arg155 | Tyr73 | b23 | E (ApcA) |
|  | Glu76 | b23 | E (ApcA) |
| Ala165 | Asp25 | b22 | B (ApcB) |
| Gly166 | Gly21 | b22 | B (ApcB) |
|  | Asp25 | b22 | B (ApcB) |
| CAN | Pro68 | b23 | E (ApcA) |
|  | Gly69 | b23 | E (ApcA) |
|  | Gly70 | b23 | E (ApcA) |
|  | Tyr73 | b23 | E (ApcA) |

**Extended Data Table 3:** Pairwise interactions within 4 Å between the OCP and the Top cylinders (Ta, left; Tb right).

| OCP<br>Residue | PBS<br>Residue | Trimer/Protein | Chain |
| --- | --- | --- | --- |
| Thr52 | Pro68 | Ta1 | A (ApcA) |
|  | Tyr73 | Ta1 | A (ApcA) |
| Ile53 | Pro68 | Ta1 | A (ApcA) |
| Ala54 | Gln60 | Ta2 | C (ApcA) |
| Ala55 | Gly69 | ta1 | A (ApcA) |
|  | Lys61 | ta2 | C (ApcA) |
|  | Pro63 | ta2 | C (ApcA) |
| Pro56 | Gln60 | ta2 | C (ApcA) |
|  | Lys61 | ta2 | C (ApcA) |
| Gly57 | Lys61 | ta2 | C (ApcA) |
|  | Arg62 | ta2 | C (ApcA) |
|  | Gln128 | ta2 | C (ApcA) |
| Ala58 | Gly69 | ta1 | A (ApcA) |
|  | Lys61 | ta2 | C (ApcA) |
|  | Arg62 | ta2 | C (ApcA) |
| Ser60 | Thr50 | ta1 | F (ApcB) |
|  | Glu54 | ta1 | F (ApcB) |
| Gln62 | Asn47 | ta1 | F (ApcB) |
|  | Thr50 | ta1 | F (ApcB) |
|  | Leu140 | ta1 | F (ApcB) |
|  | Arg131 | ta2 | C (ApcA) |
| Leu63 | Thr50 | ta1 | F (ApcB) |
|  | Glu54 | ta1 | F (ApcB) |
|  | Val136 | ta1 | F (ApcB) |
|  | Leu140 | ta1 | F (ApcB) |
| Asn66 | Ser139 | ta1 | F (ApcB) |
|  | Leu140 | ta1 | F (ApcB) |
|  | Gly142 | ta1 | F (ApcB) |
| Thr97 | Ser139 | ta1 | F (ApcB) |
| Ser100 | Lys58 | ta1 | F (ApcB) |
|  | Ser139 | ta1 | F (ApcB) |
| Trp101 | Ser139 | ta1 | F (ApcB) |
| Ser102 | Glu54 | ta1 | F (ApcB) |
| Pro103 | Arg249 | cpcg1 |  |
| Ile105 | Glu54 | ta1 | F (ApcB) |
| Glu146 | Lys234 | cpcg1 |  |
| Ser147 | Tyr73 | ta1 | A (ApcA) |
| Gly148 | Gly74 | ta1 | A (ApcA) |
|  | Glu75 | ta1 | A (ApcA) |
| Ile151 | Tyr73 | ta1 | A (ApcA) |
|  | Gly74 | ta1 | A (ApcA) |
| Thr152 | Gly74 | ta1 | A (ApcA) |
|  | Glu75 | ta1 | A (ApcA) |
|  | Glu76 | ta1 | A (ApcA) |
| Arg155 | Tyr73 | ta1 | A (ApcA) |
|  | Gly74 | ta1 | A (ApcA) |
|  | Met77 | ta1 | A (ApcA) |
| Asn314 | Leu140 | ta2 | C (ApcA) |
| Phe315 | Gly139 | ta2 | C (ApcA) |
|  | Leu140 | ta2 | C (ApcA) |
| Arg317 | Thr42 | ta2 | C (ApcA) |
| CAN | Pro68 | ta1 | A (ApcA) |
|  | Gly69 | ta1 | A (ApcA) |
|  | Gly70 | ta1 | A (ApcA) |
|  | Tyr73 | ta1 | A (ApcA) |
| Val21 | Glu75 | Tb1 | A (ApcA) |
| Thr52 | Pro68 | tb1 | A (ApcA) |
|  | Tyr73 | tb1 | A (ApcA) |
| Ile53 | Pro68 | Tb1 | A (ApcA) |
|  | Asn230 | cpcg1 |  |
| Ala54 | Gln60 | tb2 | C (ApcA) |
| Ala55 | Gly69 | tb1 | A (ApcA) |
|  | Lys61 | tb2 | C (ApcA) |
|  | Pro63 | tb2 | C (ApcA) |
| Pro56 | Lys61 | tb2 | C (ApcA) |
| Gly57 | Lys61 | tb2 | C (ApcA) |
|  | Arg62 | tb2 | C (ApcA) |
|  | Gln128 | tb2 | C (ApcA) |
| Ala58 | Gly69 | tb1 | A (ApcA) |
|  | Lys61 | tb2 | C (ApcA) |
|  | Arg62 | tb2 | C (ApcA) |
| Ser60 | Thr50 | tb1 | F (ApcB) |
|  | Glu54 | tb1 | F (ApcB) |
| Gln62 | Asn47 | tb1 | F (ApcB) |
|  | Thr50 | tb1 | F (ApcB) |
|  | Leu140 | tb1 | F (ApcB) |
|  | Arg131 | tb2 | C (ApcA) |
| Leu63 | Glu54 | tb1 | F (ApcB) |
|  | Ser139 | tb1 | F (ApcB) |
|  | Leu140 | tb1 | F (ApcB) |
| Asn66 | Ser139 | tb1 | F (ApcB) |
|  | Leu140 | tb1 | F (ApcB) |
|  | Gly142 | tb1 | F (ApcB) |
| Thr97 | Ser139 | tb1 | F (ApcB) |
| Ser100 | Lys58 | tb1 | F (ApcB) |
|  | Glu135 | tb1 | F (ApcB) |
| Trp101 | Ser139 | tb1 | F (ApcB) |
| Ser102 | Glu54 | tb1 | F (ApcB) |
| Ile105 | Glu54 | tb1 | F (ApcB) |
| Glu146 | Lys234 | cpcg1 |  |
|  | Val235 | cpcg1 |  |
| Ser147 | Tyr73 | tb1 | A (ApcA) |
| Gly148 | Gly74 | tb1 | A (ApcA) |
|  | Glu75 | tb1 | A (ApcA) |
| Ile151 | Tyr73 | tb1 | A (ApcA) |
|  | Gly74 | tb1 | A (ApcA) |
| Thr152 | Gly74 | tb1 | A (ApcA) |
|  | Glu75 | tb1 | A (ApcA) |
|  | Glu76 | tb1 | A (ApcA) |
| Arg155 | Tyr73 | tb1 | A (ApcA) |
|  | Glu76 | tb1 | A (ApcA) |
|  | Met77 | tb1 | A (ApcA) |
| Asn314 | Gly139 | tb2 | C (ApcA) |
| Phe315 | Gly139 | tb2 | C (ApcA) |
|  | Leu140 | tb2 | C (ApcA) |
| Arg317 | Thr42 | tb2 | C (ApcA) |
| CAN | Pro68 | tb1 | A (ApcA) |
|  | Gly69 | tb1 | A (ApcA) |
|  | Gly70 | tb1 | A (ApcA) |
|  | Tyr73 | tb1 | A (ApcA) |

## Supplementary Information

**Supplementary video 1: Display of the CTD displacement and carotenoid translocation in OCP**^**R**^. NTD = N-terminal domain, CTD = C-terminal domain, NTE = N-terminal extension. The carotenoid appears in sphere representation. β1 and β2 surfaces are labeled as well.

